# UNI-RNA: UNIVERSAL PRE-TRAINED MODELS REVOLUTIONIZE RNA RESEARCH

**DOI:** 10.1101/2023.07.11.548588

**Authors:** Xi Wang, Ruichu Gu, Zhiyuan Chen, Yongge Li, Xiaohong Ji, Guolin Ke, Han Wen

## Abstract

RNA molecules play a crucial role as intermediaries in diverse biological processes. Attaining a profound understanding of their function can substantially enhance our comprehension of life’s activities and facilitate drug development for numerous diseases. The advent of high-throughput sequencing technologies makes vast amounts of RNA sequence data accessible, which contains invaluable information and knowledge. However, deriving insights for further application from such an immense volume of data poses a significant challenge. Fortunately, recent advancements in pre-trained models have surfaced as a revolutionary solution for addressing such challenges owing to their exceptional ability to automatically mine and extract hidden knowledge from massive datasets. Inspired by the past successes, we developed a novel context-aware deep learning model named Uni-RNA that performs pre-training on the largest dataset of RNA sequences at the unprecedented scale to date. During this process, our model autonomously unraveled the obscured evolutionary and structural information embedded within the RNA sequences. As a result, through fine-tuning, our model achieved the state-of-the-art (SOTA) performances in a spectrum of downstream tasks, including both structural and functional predictions. Overall, Uni-RNA established a new research paradigm empowered by the large pre-trained model in the field of RNA, enabling the community to unlock the power of AI at a whole new level to significantly expedite the pace of research and foster groundbreaking discoveries.

## 1 Introduction

Ribonucleic acids (RNAs) are a group of macromolecules that perform a multitude of functions, acting not only as intermediaries in the flow of genetic information, but also as key regulators and structural components. The backbone of the central dogma contains 3 major RNA components: the messenger RNA (mRNA), ribosomal RNA (rRNA), and transfer RNA (tRNA). The mRNA conveys genetic information from DNA to protein. The rRNA forms the core of the ribosome, providing the catalytic activity of protein synthesis, while tRNA serves as an adaptor molecule that translates the nucleotide sequence of mRNA into amino acid sequences.

In recent years, the discovery of numerous non-coding RNAs (ncRNAs) has expanded our understanding of RNA functions and their impact on cellular processes [1]. Among these, microRNA (miRNA) emerged as a major class of small regulatory RNA that modulates gene expression post-transcriptionally, playing a crucial role in various physiological and pathological processes[2]. Small nuclear RNA (snRNA) is another type of ncRNA that, together with proteins, forms the spliceosome complex responsible for the splicing of pre-mRNA[3]. In addition, long non-coding RNA (lncRNA) has been found to participate in diverse cellular processes such as chromatin remodeling, transcriptional regulation, and RNA processing [4].

The study of RNA presents several challenges from both structural and functional perspectives. One of the main difficulties in studying RNA structures arises from their dynamic nature, which makes them highly mobile in physiological conditions, akin to intrinsically disordered proteins[5]. This characteristic often leads to the existence of multiple conformations, conformational exchanges, and transient interactions that are difficult to capture using traditional experimental techniques such as X-ray crystallography and nuclear magnetic resonance (NMR) spectroscopy[6]. Furthermore, unlike proteins, RNA molecules share much less sequence and structural similarities, complicating the identification of homologous RNA families and the computational prediction of secondary and tertiary structures [7].

Functionally, RNA molecules exhibit remarkable diversities, serving roles that span from catalytic activity to gene regulation, resulting in distinct mechanisms and interactions that are often challenging to investigate experimentally. For instance, the study of mRNA requires intricate techniques to analyze processes such as alternative splicing, RNA editing, and mRNA decay[8–10]. Similarly, the more versatile yet less understood long non-coding RNAs (lncRNAs) also necessitate in-depth examination to decipher their complex mechanisms[11, 12]. The inherent difficulty in performing experiments on RNA systems, particularly in vivo, may introduce substantial noise and lead to limited availability of high-quality data. These factors, combined with the vast diversity of RNA molecules and their cellular functions, highlight the pressing need for a unified and adaptable computational framework capable of systematically addressing the challenges associated with the structural and functional study of RNA.

However, conventional experimental and computational approaches encounter obstacles when it comes to comprehensively exploring the immense sequence and structural space of RNA[14–16]. The high-dimensional nature of RNA systems surpasses the capabilities of traditional research methodologies, limiting our ability to capture the complete landscape of RNA. Notably, large-scale pre-train models have exhibited remarkable efficacy in addressing high-dimensional problems in natural language processing[13, 17] and scientific fields such as physics[18], chemistry[19] and protein[20], where different ‘languages’ are used. The rationale of pre-training is leverage a large amount of data to learn general features and representations that can be fine-tuned or adapted for specific tasks more efficiently. These achievements along with the similarities between RNA and the aforementioned scenarios, suggest that by employing appropriate mathematical descriptions for extensive data and sophiscated model architectures, the pre-train approach holds great potential in the field of RNA.

Preliminary studies have been performed with certain success[21, 22], but the effectiveness of these pre-trained models were questioned. To be specific, in one recent study, the pre-train model was found to be worse than even one-hot representation in the downstream tasks[23]. Considering the limited datasets and inadequate model structures used in the previous study, we believe significant more work can be done to delineate the true boundary of pre-train schemes in the realm of RNA. Therefore, we have developed a series of context-aware deep learning models, called Uni-RNA. Based on the well-developed BERT architecture[13], advanced techniques such as rotary embedding, flash attention, and fused layernorm were integrated for optimal performance in terms of training efficiency and representational capabilities. These models performed pre-training using 1 billion RNA sequences from different species and categories (See Figure 1a). By fine-tuning Uni-RNA on a wide range of downstream tasks, including those with limited data available, related to RNA structure and functions, we have achieved SOTA results in all of them, demonstrating the extraordinary and omniscient ability of Uni-RNA (See Figure 1b). Moreover, we demonstrated the model indeed extracts useful hidden information, paved way for future in-depth applications to decipher the mysteries of life. The advent of Uni-RNA models heralds a paradigm shift in RNA science, liberating researchers from the limitation of traditional methods, unlocking substantial advancements by harnessing the comprehensive knowledge embedded within the model, and demonstrating the immense potential of large-scale pre-trained models in solving the complexities of RNA.

**Figure 1:**
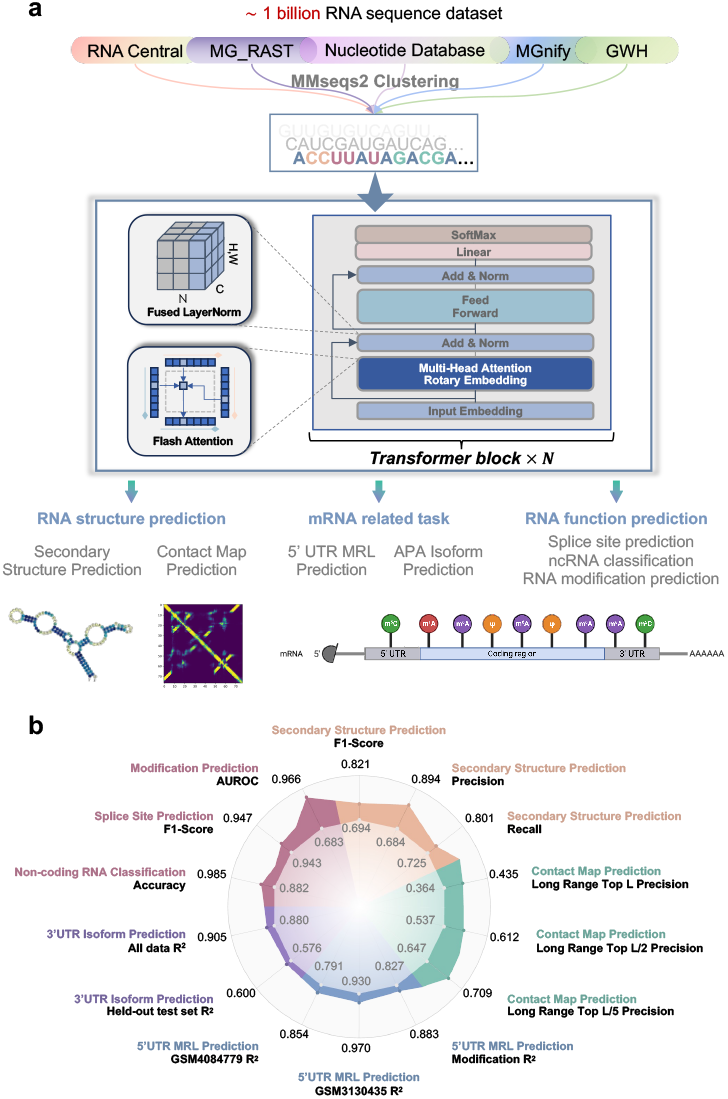
Overview of Uni-RNA models and application in RNA science. **a**. The training set for Uni-RNA models comprises sequences from five well-known RNA databases, covering both coding and non-coding RNAs. To remove sequence redundancy, MMseqs2 clustering is employed. For expediting training and scaling up the models, Flash attention and fused layernorm are utilized. The training strategy is similar with BERT[13]. After pretraining, Uni-RNA is fine-tuned on various downstream tasks, such as structure and function prediction. **b**. Uni-RNA outperforms previous SOTA and one-hot encoding methods in the downstream tasks, demonstrating superior performance.

## 2 Results

### 2.1 Uni-RNA extract hidden structural information

In the field of RNA science, accurately predicting RNA structures, including both secondary and tertiary structures, is of paramount importance. High-precision predictions of RNA structures form the basis for understanding RNA functions and revealing the complex interactions between RNA and small-molecule ligands or other bio-macromolecules. However, the expenses and difficulties in precisely characterizing RNA structures by conventional web-lab experiments hindered the accessibility to extensive RNA structure data. To tackle these problems, numerous computational methods have been proposed and extensively investigated[24–30]. In the following section, we will investigate the potential of Uni-RNA models in the prediction of RNA structures and demonstrate its superior performance.

### RNA secondary structure prediction improved by Uni-RNA models

Accurate prediction of RNA secondary structure carries broad implications, including the understanding of RNA folding mechanisms, dynamics of RNAprotein interactions, and the development of RNA-based therapeutics. Here, we conducted experiments on established benchmarks with the same datasets from E2Efold[31], UFold[32], and RNA-FM[33]. By directly fine-tuning on Uni-RNA(see **Methods**), several experiments were conducted to evaluate the performance of Uni-RNA presented in Table 1 and Figure 2a. Benchmarks of other methods were directly adopted from RNA-FM paper[33]. Considering the imbalance of classes, we chose macro averages of precision score, recall score, and F1-score as metrics to better evaluate model performance, same as previous work[33]. The Uni-RNA demonstrated outstanding performance compared to all other methods across all metrics[34–43]. Notably, in comparison to the previous SOTA model RNA-FM, Uni-RNA exhibited substantial improvements, surpassing RNA-FM by **18.3%** in F1-score, **10.4%** in recall score, and **30.7%** in precision score.

**Table 1:** Uni-RNA based RNA secondary structure prediction performance. Uni-RNA extracts latent information from billions of RNA sequences, which enables highly accurate RNA secondary structure prediction. Our model outperforms the other 13 methods with respect to all three metrics. The benchmarks of other methods were adopted from RNA-FM paper[33]

| Methods | Metric |  |  |
| --- | --- | --- | --- |
|  | Precision | Recall | F1-Score |
| Uni-RNA * | **0.894** | **0.801** | **0.821** |
| RNA-FM | 0.684 | 0.725 | 0.694 |
| UFold | 0.607 | 0.741 | 0.654 |
| E2Efold | 0.140 | 0.129 | 0.130 |
| LinearFold | 0.561 | 0.581 | 0.550 |
| Mfold | 0.501 | 0.627 | 0.538 |
| RNAstructure | 0.494 | 0.622 | 0.533 |
| RNAfold | 0.494 | 0.631 | 0.536 |
| CONTRAFold | 0.528 | 0.655 | 0.567 |
| SPOT-RNA | 0.594 | 0.693 | 0.619 |
| RNAsoft | 0.497 | 0.626 | 0.535 |
| MXfold2 | 0.519 | 0.646 | 0.558 |
| Contextfold | 0.529 | 0.607 | 0.546 |
| Eternafold | 0.516 | 0.666 | 0.563 |

\* Uni-RNA L24 model.

**Figure 2:**
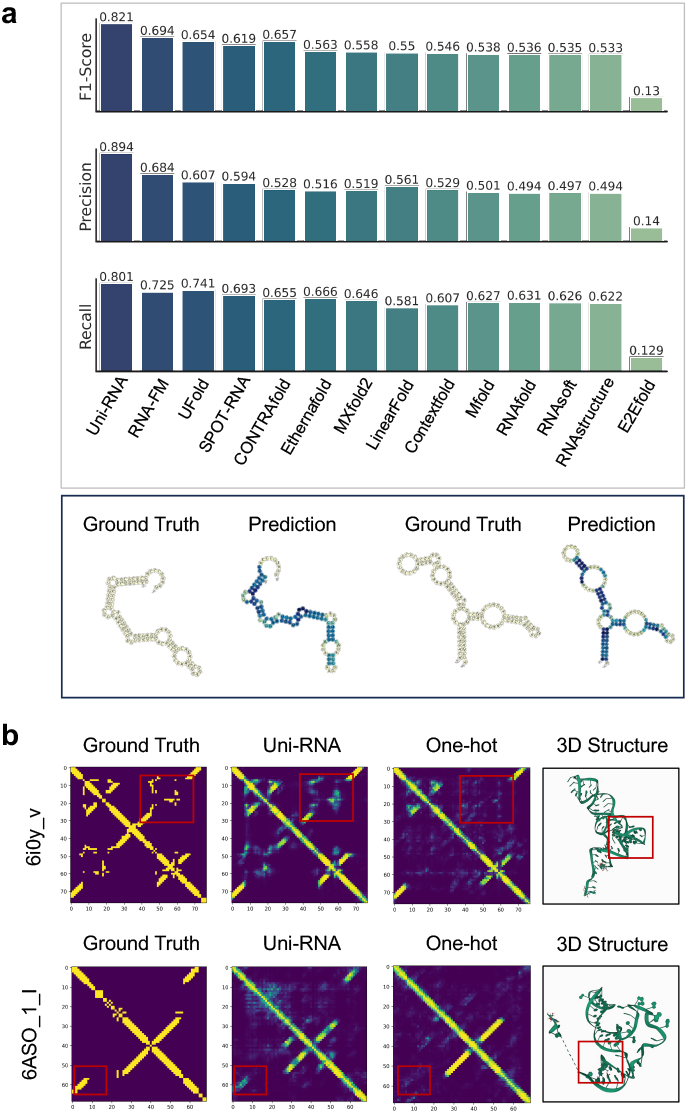
Uni-RNA’s application on RNA structure prediction tasks. We evaluated on commonly used RNA secondary structure dataset: bpRNA-1m and PDB data. **a**. The bar chart compares the performance of different methods on the test dataset, with F1-score, precision, and recall as our metrics. Uni-RNA model outperforms all previous methods on all three metrics. **b**. We compared the contact maps predicted by Uni-RNA and the structures predicted by the classical one-hot encoding method. In contrast to the commonly used one-hot encoding, the predicted structures obtained by the Uni-RNA model better capture long-range dependency relationships.

### Uni-RNA benefits RNA tertiary structure prediction

The tertiary structure of RNA is essential for understanding the mechanisms involved in RNA-mediated processes, such as gene expression, protein synthesis and cellular regulation[44]. Unlike the secondary structure, which primarily reveals local base pair interactions, the contact map provides additional information in the three-dimensional space[45]. While the secondary structure can impose strict constraints on structure modeling, the contact map offers more loose yet global constraints that can be critical for accurate modeling. Here we represented contact map as a two-dimensional matrix where each point corresponds to the pairwise inter-nucleotide distance in the tertiary structure. The distances were measured between the center of mass of any two bases, and a threshold of 8 Å was set.

To conduct a comprehensive evaluation of Uni-RNA’s performance in predicting RNA contact maps, we constructed a dataset based on 658 curated high quality experimental RNA structures. Table 2 illustrates the long-range top precision of different methods on the test set. To enable a fair comparison, we employed the same network for contact map prediction but utilized different representation methods. Uni-RNA demonstrated a significant improvement in prediction accuracy on the test set compared to one-hot encoding. Particularly, in several test cases, we observed that Uni-RNA can capture the off-diagonal long range contacts where one-hot representation cannot (Figure 2b). This superior performance can be attributed to the hidden structural and functional information extracted by the pre-trained model.

**Table 2:** Uni-RNA performance on RNA contact map prediction. The structure information extracted by Uni-RNA models significantly enhances the prediction accuracy, surpassing the commonly used one-hot encoding by over 20%. The top precision metric calculates the percentage of correctly predicted contacts among the top L, L/2, and L/5 predicted contacts compared to the true contacts. The predicted contacts are ranked based on their confidence scores, and the top L, L/2, and L/5 contacts are selected for evaluation. Furthermore, long-range evaluation refers to the assessment of contacts where the sequence separation between two residues exceeds 24. It focuses on evaluating the accuracy of long-range contacts specifically.

| Methods | Long-Range Top Precision |  |  |
| --- | --- | --- | --- |
|  | L | L/2 | L/5 |
| Uni-RNA * | **0.435** | **0.612** | **0.709** |
| One-hot encoding | 0.364 | 0.537 | 0.647 |

\* Uni-RNA L16 model.

### 2.2 Uni-RNA facilitates the development of mRNA therapeutics

In the past decade, there has been a rising trend in the research and clinical development to manipulate mRNA to achieve diverse physiological functions[46, 47]. In this section, we will explore the potential applications of the Uni-RNA model in mRNA related tasks. Based on the evolutionary information learned from extensive RNA data, we aimed to employ the Uni-RNA model to investigate some of the most critical dark corners within the untranslated regions(UTRs), besides the coding regions. The results suggest our models present novel opportunities for advancing research in mRNA-based therapy.

### Ribosome load prediction based on 5’UTR sequence

The sequence of 5’UTR plays a pivotal role in governing translation efficiency. Mean Ribosome Load (MRL) is used to reflect the translational activities and protein synthesis levels of mRNA under specific conditions. Precise prediction of ribosome load capacities based on the 5’UTR sequence offers invaluable guidance for mRNA sequence design towards optimal protein expression outcomes, especially to go beyond the framework of existing 5’UTR templates towards novel sequence designing. We conduct experiments on benchmarks used in previous work[48]. This work employed Massively Parallel Reporter Assays (MPRAs) to construct a library comprising 280,000 gene sequences and calculated the corresponding ribosome loading. Based on those data, we fine-tuned the Uni-RNA to predict the MRL from the 5’UTR sequences (see **Methods**). Besides, we also utilized the datasets employed by RNA-FM[33], which include random data (Random 7600) and human data (Human 7600).

To further validate the robustness of our approach, we conducted additional experiments involving the evaluation of different coding sequences (CDS) replacements including EGFP and mCherry and chemically modified RNA sequences. We conducted tests on these sets of experimental data to evaluate the performance of Uni-RNA by *R*^2^(See Figure 3a). In different experiments, Uni-RNA demonstrates potent representational capabilities and achieves better results across all tasks shown in Table 3.

**Table 3:** Uni-RNA performance (*R*^2^) on 5’UTR sequence MRL prediction. EGFP and mCherry are two different methods of replacing coding sequences (CDS), based on a library containing 280,000 sequences. Ψ and *m*_1_Ψ represent two types of RNA modification. Rep1 and Rep2 are evaluations performed on two independent polysome profiling experimental data. Furthermore, we conducted experiments to compare our approach with RNA-FM, using the same dataset Random 7600 and Human 7600. Uni-RNA consistently demonstrated better prediction accuracy and robust performance in different datasets, CDS replacing methods, and modification types.

| Model | EGFP |  |  |  |  |  | mCherry |  | Human dataset |  |
| --- | --- | --- | --- | --- | --- | --- | --- | --- | --- | --- |
| | Unmodified | | $\Psi$ | | $m_1\Psi$ | | Rep1 | Rep2 | Random 7600 | Human 7600 |
|  | Rep1 | Rep2 | Rep1 | Rep2 | Rep1 | Rep2 |  |  |  |  |
| UniRNA* | **0.96** | **0.92** | **0.82** | **0.92** | **0.80** | **0.88** | **0.87** | **0.86** | **0.91** | **0.85** |
| Optimus 5-Prime | 0.91 | 0.87 | 0.78 | 0.82 | 0.77 | 0.81 | 0.85 | 0.84 | 0.84 | 0.78 |
| RNA-FM | - | - | - | - | - | - | - | - | 0.85 | 0.79 |

\* Uni-RNA L16 model.

**Figure 3:**
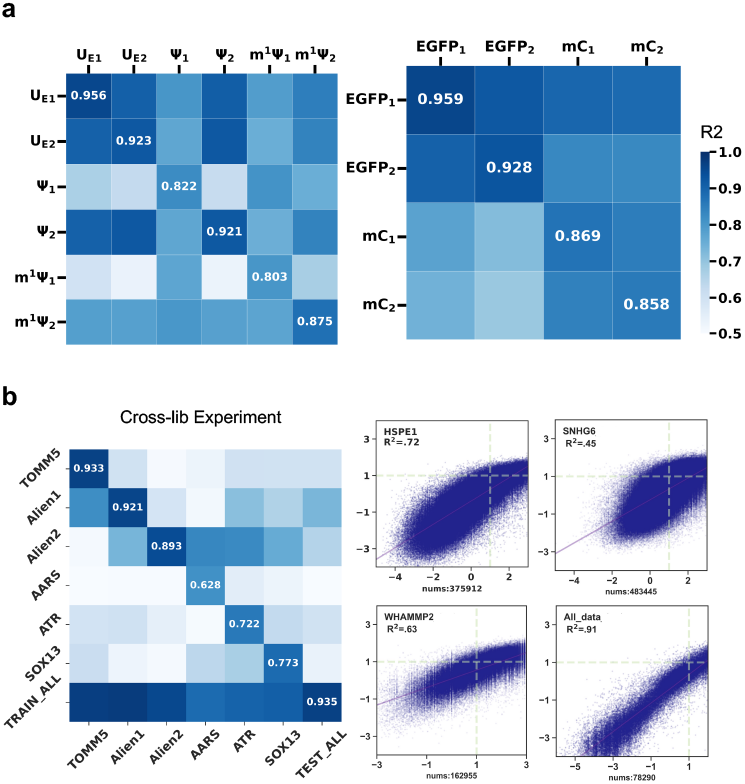
Uni-RNA preformance on 5’ UTRs ribosome load and APA isoform type prediction. **a**. Model performance (*R*^2^) with training and testing on different 5’ UTR MRL prediction datasets. *U*_*E*1_, *U*_*E*2_, Ψ_1_, Ψ_2_, *m*^1^Ψ_1_, *m*^1^Ψ_2_ represent different RNA modifications. *EGFP*_1_, *EGFP*_2_, *mC*_1_, *mC*_2_ represent different CDS sequences combined with the same collection of 5’UTR sequences. **b**. Model performance (*R*^2^) with training and testing on different APA isoform prediction datasets. HSPE1, SNHG6, and WHAMMP2 represent three held out test sets. All-data stands for randomly split test set.

#### Isoform percentage prediction based on 3’UTR sequence

Alternative polyadenylation (APA) is a widespread phenomenon observed in various organisms, playing a crucial role in gene expression regulation and cellular processes. It facilitates the generation of mRNA isoforms with different characteristics, thereby enabling precise modulation of gene expression levels and diversification of protein functions. Here, we followed APARENT[49] to train a model to predict the proximal APA isoform ratio for each variant 3’UTR sequence. To assess the generalization ability of the models, we conducted experiments on different datasets (see **Methods**), including 9 libraries spliting into train, validation and test and three held-out datasets (HSPE1, SNHG6, WHAMMP2) for addtional stand-alone tests (See Figure 3b right). In comparison to the previous SOTA method, Uni-RNA consistently demonstrated superior performance as shown in Table 4.

**Table 4:** Uni-RNA performance (*R*^2^) on isoform percentage prediction based on 3’UTR sequence. The isoform percentage prediction task utilizes the 3’UTR sequence embeddings obtained through Uni-RNA extraction to predict isoform abundance. The training was performed using a combined dataset comprising approximately 2.5 million sequences from 9 libraries. To valid the robustness of our model, we did further cross-lib tests. The Total metric represents the *R*^2^ on the all test sets combined. Mean and Min represent the average and minimum values of the 9 libs, respectively, represented by the diagonal entries in the cross-library test (Figure 3b left). The Uni-RNA model outperforms previous SOTA on both the combined data and the individual libraries. Moreover, Uni-RNA exhibited improved generalization capabilities on 3 independent test libraries.

| Model | 9 libs |  |  | 3 held-out libs |  |  |
| --- | --- | --- | --- | --- | --- | --- |
|  | Total | Mean | Min | HSPE1 | SNHG6 | WHAMMP2 |
| Uni-RNA * | **0.91** | **0.81** | **0.63** | **0.72** | **0.45** | **0.63** |
| APARENT | 0.88 | 0.68 | 0.50 | 0.69 | 0.45 | 0.59 |

\* Uni-RNA L16 model.

### 2.3 Uni-RNA reveals potential relationships between sequence and function

Predicting RNA functional attributes from its sequence holds special meaning in the field of RNA. Through correct predictions of these attributes, scientists can obtain deep understanding of diverse biological mechanisms and effectively alter and create RNA molecules for various purposes. Nevertheless, the resource-demanding and elaborate experimental methodologies have difficulties in conducting such functional studies on a large scale, while the existing computational approaches straggled to decode the complicated nature of RNA with limited data. Encouragingly, Uni-RNA leverages the strength of pre-training to augment sequence representation, consequently allowing for improved performance in a wide range of downstream tasks without additional datasets.

#### Cross-species splice site prediction

Splicing, a regulatory mechanism of gene expression, involves the removal of introns and the joining of exons during precursor mRNA transcription, to produce the mature mRNA molecules. Accurate identification of splice sites is critical for proper splicing, as it directly influences the integrity of subsequent protein translation. In this study, we used Uni-RNA models to predict splice sites on mRNA sequences. To comprehensively evaluate the model’s performance, we conducted comparative analyses with several established methods, including DNABERT[50], Spliceator[51], SpliceFinder[52], DSSP[53], MaxEntScan[54], NNSplice[55], and SpliceBERT[56]. F1-score was used as the evaluation metric, and the results are presented in Table 5. The SOTA performance of our model demonstrates its remarkable generalization capabilities across multiple species. This can be attributed to the power of pre-training on one billion high-quality RNA sequences from various sources.

**Table 5:** Uni-RNA facilitates cross-species prediction of splice sites. Efficient prediction of splice sites will be beneficial in uncovering the intricacies of gene regulation and the molecular mechanisms of diseases. The Uni-RNA model achieves SOTA performance on test sets from four different species, further highlighting its strong generalization capabilities.

| Methods | Species |  |  |  |
| --- | --- | --- | --- | --- |
|  | zebrafish | fruit fly | worm | arabidopsis |
| Uni-RNA * | **0.9635** | **0.9498** | **0.9394** | **0.9362** |
| SpliceBERT | 0.9568 | 0.9461 | 0.9343 | 0.9361 |
| DNABERT | 0.9505 | 0.9307 | 0.9094 | 0.9094 |
| Spliceator | 0.935 | 0.929 | 0.916 | 0.929 |
| DSSP | 0.94 | 0.927 | 0.86 | 0.852 |
| MaxEntScan | 0.899 | 0.91 | 0.884 | 0.896 |
| SpliceFinder | 0.901 | 0.857 | 0.794 | 0.793 |
| NNSplice | 0.743 | 0.762 | 0.635 | 0.648 |

\* Uni-RNA L16 model.

#### Classification of ncRNA functional families

The ncRNAs represent a class of transcripts that do not encode proteins but exert critical regulatory functions across diverse biological processes and diseases. Despite their significance, the understanding of their biological functions remains highly incomplete. While certain existing methods show promise in predicting the function of ncRNAs based on secondary structures, here we employed sequence-based approaches based on Uni-RNA to offer a computationally efficient solution with exceptional representation capabilities for different types of ncRNAs. We followed the work from ref[59] and evaluated the model performance on ncRNA family classification tasks. Additionally, we examined the model’s accuracy when confronted with uncertainty regarding the start and end positions of ncRNA sequences. This uncertainty can arise from the noises introduced during the next-generation sequencing process, or the natural mutations. To simulate this scenario, we introduced varying amounts of boundary noises to each sequence. The boundary noises consisted of a random number of nucleotides added to both ends of the sequence, while maintaining the same single-nucleotide and di-nucleotide frequencies to the original sequence. We explored noise lengths ranging from 0% to 200% of the original sequence length.

Compared with existing methods, our model achieved SOTA performance on both noise-free and noisy data (see Table 6). Furthermore, the classification performance of the Uni-RNA model remained stable across different levels of boundary noise (see Figure 4). These results further validate the robust and excellent performance of the Uni-RNA model, especially in the sense of extracting evolutionary information.

**Table 6:** Highly accurate prediction of short non-coding RNA functions by utilizing Uni-RNA models. The Uni-RNA model demonstrated SOTA performance compared to other methods at various levels of boundary noise. The 3-mer, 2-mer, and 1-mer methods are different approaches for sequence representation. Hibert [57], Morton [58], and Snake [59] refer to three distinct types of 2D space-filling curves. EdeN[60] and nRC[61] are two deep learning-based architectures.

| Methods | Accuracy |  |
| --- | --- | --- |
|  | 0% boundary noise | 200% boundary noise |
| Uni-RNA * | **0.98** | **0.98** |
| 3-mer | 0.89 | 0.84 |
| 2-mer | 0.88 | 0.84 |
| 1-mer | 0.87 | 0.81 |
| Snake | 0.82 | 0.70 |
| Morton | 0.82 | 0.68 |
| Hibert | 0.83 | 0.67 |
| EdeN | 0.88 | 0.68 |
| nRC | 0.90 | 0.64 |

\* Uni-RNA L16 model.

**Table 7:** Uni-RNA improves the accurate prediction of multiple RNA modification sites. Uni-RNA demonstrates excellent predictive performance across 12 artificial and natural RNA modification categories, exhibiting significant performance improvement compared to the commonly used one-hot encoding method.

| Methods | Species |  |  |  |  |  |  |  |  |  |  |  |
| --- | --- | --- | --- | --- | --- | --- | --- | --- | --- | --- | --- | --- |
| | *Am* | *Cm* | *Gm* | *Tm* | $m^1A$ | $m^5C$ | $m^5U$ | $m^6A$ | $m^6Am$ | $m^7G$ | $\Psi$ | I |
| Uni-RNA* | **0.929** | **0.968** | **0.986** | **0.959** | **0.954** | **0.976** | **0.958** | **0.994** | **0.978** | **0.956** | **0.942** | **0.993** |
| One-hot encoding | 0.648 | 0.617 | 0.571 | 0.658 | 0.699 | 0.681 | 0.752 | 0.689 | 0.621 | 0.592 | 0.687 | 0.980 |

\* Uni-RNA L16 model.

**Figure 4:**
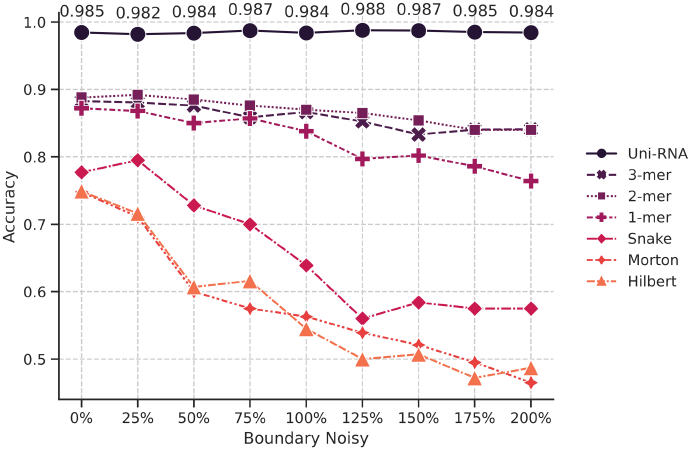
Uni-RNA models empower RNA function prediction. The classification accuracy of different methods was evaluated by introducing boundary noise at various levels. As the level of boundary noise increased, Uni-RNA demonstrated consistent and robust performance.

#### Prediction of RNA modification sites

Post-transcriptional RNA modifications play a crucial role in the regulation of the epi-transcriptome across all types of RNA. Accurate identification of RNA modification sites is of paramount importance for elucidating the functional implications and regulatory mechanisms underlying RNA molecules. In this study, we fine-tuned the Uni-RNA model to predict RNA modification sites. Predicting the full-length RNA sequence is of greater biological significance, yet prior approaches, constrained by their limited model capacity, struggled to effectively handle longer RNA sequences. Uni-RNA, however, is capable of accommodating RNA sequences of varying lengths due to its advanced model design, enabling direct prediction of modifications across full-length sequences. This advancement provides a more comprehensive understanding of RNA modifications and their functional implications. As to the evaluation metric, consistent with the previous studies[62], we computed the AUROC using positive samples and their corresponding negative samples for each modification. The results are summarized in Table 5. Uni-RNA outperformed one-hot encoding in all 12 modification types, demonstrating superior performance.

## 3 Discussion

In summary, we have developed Uni-RNA, a large-scale pre-training model for RNA sequences. It has been trained on one billion RNA sequences and demonstrated SOTA performance in various downstream tasks related to RNA structure and function predictions. We observed that as the training data and model parameters increased, the performance of the Uni-RNA model in downstream tasks also improved, which aligns with the observation that large-scale pretraining enables the training of deeper models with greater predictive potential. However, when the model parameters exceeded 400M, the performance in downstream tasks reached plateau with the current architectures and datasets. The accurate prediction of RNA structures by Uni-RNA establishes the physical foundation for studying RNA functions. Furthermore, its ability to predict mRNA-related tasks expedites the research and development of mRNA therapies. It’s worth mentioning the Uni-RNA model exhibited extraordinary classification performance for different categories of RNA, including pre-mRNA and non-coding RNA (ncRNA), indicating the Uni-RNA begins to understand the nature of RNA molecules facilitating different biological functions. Consequently, the direct predictions of RNA function from sequences were achieved at high accuracies, which can help researchers in exploring novel RNA mechanisms. With future studies, we envision the Uni-RNA can help enabling efficient RNA therapy development, including ASO[63], SiRNA[64], Aptamer[65] and RNA targeting small molecules[66], given more corresponding experimental data and further training.

In the foreseeable future, the RNA research will inevitable play a more and more important role towards decrypting the enigma of life. It is anticipated as the fast advancement of biotechniques, a surge of data in various form will continue to emerge at an exponential rate, which calls for sophisticated and automated frameworks to gather and process to convey valuable information and knowledge. We believe the Uni-RNA stands as a groundbreaking prototype of such framework, as our investigation demonstrates its universal capabilities. Moreover, the large language models (LLM) like GPT-4 have profoundly changed our life, including the scientific research, since the majority of our information and knowledge is carried by natural languages. However, there still exists a huge gap between LLMs and biological science, because the biological science speaks different languages mainly in amino acid or nucleotide sequences. We believe the Uni-RNA can serve as the nexus to bridge between the biological context and our knowledge, through techniques such as cross modality feature alignments, to ultimately help us unravel the laws of the RNA universe.

Applications and codes will be provided via Bohrium^®^ Apps and Bohrium^®^ notebooks (https://bohrium.dp.tech/).

## 4 Methods

### 4.1 Training Dataset

The development of robust and efficient pre-training models hinges upon the availability of large-scale and high-quality data. In this study, we present a comprehensive RNA sequence database with sheer scale and high quality, serving as the cornerstone for the success of our model.

#### Database Construction and Data Collection

To construct this database, we embarked on an exhaustive data collection process, aggregating RNA-related nucleic acid databases from various sources, similar to one recent seminal work[67]. We incorporated non-coding RNA sequences from RNAcentral[68], nucleic acid data from NCBI’s database (nt)[69], and genomic data from repositories such as Genome Warehouse (GWH).[70] This extensive integration ensures a diverse and representative collection of RNA sequences, thereby maximizing the utility and generalizability of our database.

#### Data Alignment and Refinement

In order to maintain consistency and facilitate downstream analyses, we aligned all gathered sequences to a standardized DNA alphabet. We then performed length-based statistical analyses on the aligned sequences. To ensure optimal data quality and manageability, we excluded sequences exceeding 4096 nucleotides in length. The remaining sequences were systematically classified according to their respective origins, enabling a more granular understanding of the data.

To further refine our database and reduce redundancy, we employed the mmseqs2 clustering algorithm [71]. This approach allowed us to distill our dataset into a collection of high-quality and unique RNA sequences. Following rigorous quality control measures, we curated approximately 1 billion valid data points to formulate the database.

### 4.2 Model

In this work, we introduce a transformer-based architecture to unravel the underlying patterns of RNA sequences. We devise a masked nucleic acid modeling framework to pre-train the architecture, which enables the model to learn robust representations of the intricate biological structure.

#### Sequence Tokenization

In this study, we encoded each nucleotide (A, G, C, T/U) as a token to facilitate the extraction of hidden states and attention weights for individual nucleotides. We used “N” to represent other rare bases. Considering that some RNA sequences were not converted to the RNA alphabet, during training, we transformed all “U” bases to “T,” which aligns with the common practice of converting RNA to cDNA during actual RNA sequencing. Additionally, a “[CLS]” (classification) token and a “[SEP]” (separator) token were added at the beginning and the end of the input sequence to Uni-RNA models.

#### Model Architecture

Analogous to the ESM, we employ a BERT-style encoder-only transformer to model complex dependencies among input features. As we scale the UniRNA model, we judiciously modify the number of layers, attention heads, hidden size, and feed-forward network (FFN) hidden size Table 8 illustrates the nuances of our proposed architecture.

**Table 8:** Model architecture parameters of different Uni-RNA models. To cater to the requirements of various downstream tasks, we have trained a range of Uni-RNA models with different sizes. The table provides a detailed list of the number of layers, embedding size, model parameter size, attention heads, and training set sequence count for the Transformer models. Notably, the hidden size of FFN is three times of the embedding size.

| Models | Parameters |  |  |  |  |
| --- | --- | --- | --- | --- | --- |
|  | Number of layers | Embedding size | Attention heads | Params | Training data |
| Uni-RNA-L8 | 8 | 512 | 8 | 25M | 100M |
| Uni-RNA-L12 | 12 | 768 | 12 | 85M | 100M |
| Uni-RNA-L16 | 16 | 1024 | 16 | 169M | 500M |
| Uni-RNA-L24 | 24 | 1280 | 20 | 400M | 500M |

A transformer encoder layer comprises two primary components: multi-head attention and feed-forward network (FFN), adept at capturing both local and global contextual information. The multi-head attention mechanism is responsible for capturing different aspects of the input by computing the scaled-dot product between query *q*∈ R^*n×d*^*h*, key *k*∈R^*n×d*^*h*, and value *v*∈ R^*n×d*^*h* representations, where *n* represents the length of a sequence and *d*_*h*_ represents the hidden size of multi-head attention. The FFN consists of two fully connected layers, which enable the model to introduce complex non-linearity to the input. Owing to the position-agnostic nature of the transformer architecture, we integrate rotary embedding[72] to encapsulate positional information into the model.

These modules are interconnected via residual connections and layer normalization operations to facilitate seamless information flow and gradient propagation. Equation 5 formally delineates the computational process of the transformer encoder layer.

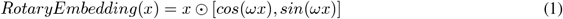

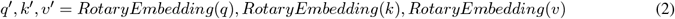

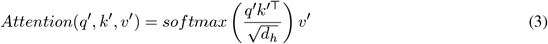

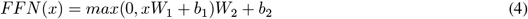

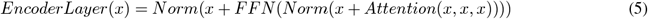

#### Implementation Details

In the pursuit of enhancing the efficacy of our proposed model, we have employed the IO-aware Flash Attention mechanism[73] in place of the traditional MultiHeadAttention. The conventional MultiHeadAttention, which involves multiple dot products, often suffers from reduced efficiency when dealing with elongated sequences. Flash Attention, on the other hand, leverages the concept of tiling to minimize the number of memory reads and writes occurring between the GPU high bandwidth memory (HBM) and the GPU on-chip SRAM. This approach enables us to attain a substantial five-fold acceleration in the processing speed, thereby allowing efficient training.

#### Training Details

The network is trained using a comprehensive pipeline that incorporates several essential components, such as a robust learning rate scheduling strategy and gradient clipping mechanisms. These elements work in synergy to promote stable convergence and mitigate potential issues pertaining to exploding or vanishing gradients. We trained our proposed network on 128 A100. The detailed training parameters for models with different scales is show on Table 9.

**Table 9:** Training parameters for Uni-RNA.

| Models | Training Paramters |  |  |  |
| --- | --- | --- | --- | --- |
|  | Peak learning rate | Step | Dropout | Weight Decay |
| Uni-RNA-L8 | 1e-4 | 400K | 0.1 | 0.00 |
| Uni-RNA-L12 | 1e-4 | 400K | 0.1 | 0.00 |
| Uni-RNA-L16 | 1e-4 | 400K | 0.1 | 0.01 |
| Uni-RNA-L24 | 1e-5 | 300K | 0.1 | 0.01 |

### 4.3 Downstream Task Datasets

#### RNA Secondary Structure Dataset

The benchmark dataset is built according to Ufold[32] and RNA-FM[33] from two sources: (1) The RNAStralign dataset[74], encompassing 37,149 structures across eight RNA types, represents one of the most extensive assemblages of RNA structures within the discipline. (2) bpRNA-1m dataset[75], which has been preprocessed by eliminating sequence similarity through an 80% sequence-identity cut-off and constraining the maximum sequence length to less than 500. The bpRNA dataset was partitioned randomly into three distinct datasets: one for training purposes, another for validation (referred to as VL0), and a third for testing (referred to as TS0). To enhance the training process, the RNAStralign dataset was combined with the training set from the bpRNA dataset, forming the training set denoted as TR0. Considering the issue of redundancy, we chose not to use the ArchiveII dataset as our test set.

#### RNA Contact Map Dataset

Initially, all RNA-containing structures with a resolution of 4 ≤ Å were downloaded from the PDB[76] website as of March 16, 2022. To ensure data quality, we removed non-RNA fragments and structures affected by proteins, small molecules, or DNA. After these preprocess steps, redundant structures were removed using CD-HIT-EST[77]. The dataset was then randomly divided into training, validation, and test sets in an 8:1:1 ratio. Subsequently, CD-HIT-EST was applied to the validation and test sets to remove redundancy based on a similarity threshold of 0.8. This process resulted in three datasets: TR0 (526), VL0 (65), and TS0 (67).

#### 5’UTR Sequence Function Prediction Dataset

The benchmark dataset for predicting MRL based on the 5’UTR is derived from massively parallel reporter assays conducted by Paul et al[48]. The experiments document mRNA sequences along with their corresponding mean ribosomal loads. The original sequences include: (1) A library consisting of 280,000 gene sequences (GSM4084779) containing a randomized 5’ UTR and a constant region comprising the coding sequence for enhanced green fluorescent protein (eGFP) and a 3’ UTR. (2) A separate 5’ UTR mRNA library with the CDS for eGFP replaced by the coding sequence for mCherry. (3) mRNA sequences modified with pseudouridine(*ψ*) and 1-methylpseudouridine(*m*^1^*ψ*). (4) A total of 7,600 sequences are randomly sampled from the RNA-FM dataset[33], which consists of 83,919 5’UTRs of 75 different lengths (GSM3130435). These sequences are evenly distributed across the length categories and used as the Random 7600 test set. The remaining sequences is adopted for train set. The Human 7600 dataset consists of 7,600 held-out real human 5’UTRs, and its length distribution aligns with that of the Random 7600 dataset.

#### Alternative Polyadenylation Dataset

The benchmark dataset for predicting the proximal isoform ratio is obtained from the extensive study conducted by Bogard et al.[49]. In their research, they systematically constructed and transiently expressed minigene libraries comprising over 3 million distinct UTR sequences. The isoform and cleavage data were subsequently extracted from the expressed RNA. We utilize a large-scale random 3’UTR libraries as the dataset for this task. The dataset comprised 12 distinct libraries of different sizes, with 9 libraries allocated for training and the remaining 3 libraries (HSPE1, SNHG6, WHAMMP2) held in reserve. Specifically, during the training phase, 95% of the data from these 9 libraries (∼2.5M sequences) was utilized, with 2% (∼50,000 sequences) for validation, and 3% (∼80,000 sequences) set aside for testing. In the cross-lib test, we followed the methodology described in the original study to process the library, got six merged libraries and conduct followed experiments

#### Splice Site Prediction Dataset

The training dataset for our model is obtained from Spliceator[51], which is built upon the G_3_PO+ benchmark. This dataset consists of curated, confirmed error-free splice sites from over 100 eukaryotic species. Additionally, Spliceator provides five independent test datasets from different organisms, including vertebrates (human and zebrafish), invertebrates (fruit fly and worm), and plants (arabidopsis). Since the training data already includes human splice sites, we focused our evaluation on the four non-human test datasets to assess the model’s performance in cross-species splice site prediction.

#### Non-coding RNA Function Analysis Dataset

The benchmark datasets were constructed based on the work reported by Cerulo et al.[59]. Starting from the original Rfam dataset, this work deleted families whose clustering was highly correlated with sequence length and families with too few RNA sequences to ensure data quality. Additionally, to evaluate the algorithm’s generalization capability, each Rfam class in the dataset was randomly divided into three subsets: training (84%), validation (8%), and testing (8%). To mitigate potential bias arising from an over-representation of highly similar homologous sequences, the protocol ensured that all sequences in the validation and test sets for each class exhibited a normalized Hamming distance of less than 0.50 with any sequence in the training set.

#### RNA Modification Sites Prediction Dataset

The benchmark dataset for RNA modification is derived from the study conducted by Song et al.[62] The investigation involved the acquisition of a comprehensive collection of 20 epi-transcriptome profiles, generated using 15 distinct base-resolution technologies, encompassing 12 diverse types of RNA modifications (*Am, Cm, Gm, Tm, m*^1^*A, m*^5^*C, m*^5^*U, m*^6^*A, m*^6^*Am, m*^7^*G*, Ψ, *I*). These profiles encompass a wide spectrum of RNA modification data, thereby providing extensive coverage. Negative sites within the dataset were randomly selected from the unmodified bases present in the same transcript that harbors the positive sites.

### 4.4 Downstream Tasks

#### Applying Uni-RNA in Various Downstream Tasks

After pretraining, we obtain the Uni-RNA model that encodes RNA’s latent structural and functional information. This model can be integrated into various downstream applications through two approaches: linear probing and fine-tuning. Linear probing is a common technique in deep learning that utilizes pretrained models for downstream tasks. During this process, the weights of the pretrained model are typically frozen, enabling the linear layer to learn task-specific features and make predictions. It offers a straightforward and efficient approach to transferring knowledge from pretrained models to new tasks. Fine-tuning is another approach used to transfer knowledge from a pretrained model to specific tasks. Unlike the linear probing strategy that freezes the entire model, fine-tuning enables adjustment of the pretrained model’s weights to better adapt to the new task. By updating the parameters through a smaller number of training iterations, the model can learn task-specific features and enhance its performance on the target task. In comparison to linear probing, fine-tuning consistently yields better performance on downstream tasks.

Uni-RNA models have been trained on billions of high-quality RNA sequences, extracting latent structural and functional patterns. The information embedded in the Uni-RNA model is expected to benefit a range of RNA-related tasks, such as structure prediction and biological function prediction.

#### RNA High-order Structure Map Prediction

RNA high-order structures, such as RNA secondary structure and RNA contact map, can be represented by a two-dimensional matrix. The secondary structure reflects hydrogen bonds in the primary sequence, while the contact map focuses on pairwise tertiary inter-nucleotide interactions. We employ a simple strategy for structure map prediction. The representation from Uni-RNA will be outer concatenated into 2D feature matrix. Subsequently, the feature map is passed through two layers of ResNet’s Bottleneck block. The task can be viewed as pixel-level classification. In the case of secondary structure prediction, each base-pair point in the 2D matrix is assigned one of two classes: 1 for paired bases and 0 for unpaired bases. Conversely, for side-chain contact map prediction, the distances of pairwise inter-nucleotides are divided into 20 bins. Two classes are used for distances exceeding 20Å and smaller than 2Å, while for distances ranging from 2Å to 20Å, 18 classes are utilized. In total, there are 20 classes for each pairwise point.

#### 5’UTR Sequence Function Prediction

For predicting the Mean Ribosome Load (MRL) value of 5UTR sequences, we take the entire sequence as input. First, we input the sequence into Uni-RNA to obtain a sequence representation. For the Uni-RNA-L16 model, we obtain a 1024 dims embedding. Then we apply a linear projection to reduce the embedding dimension from 1024 to 32. The dimensionality-reduced features are then fed into 6 layers 1D residual network and a linear layer to obtain the predicted MRL. The 1D residual network was constructed as two convolutional layers with 3 kernel sizes followed by an InstanceNorm layer and an ELU activation at each layer. The flattened convolution output is passed to a linear layer of with 0.2x dropout to get the final prediction.

#### Alternative Polyadenylation Prediction

For alternative polyadenylation prediction task, we utilize Uni-RNA-L16 model to accurately predict the isoform percentage. Our prediction pipeline aligns with the model APARENT proposed by Bogard et al.[49], with the difference being the replacement of one-hot inputs with embedding obtained by UniRNA model to demonstrate the extraction capability of our model for the biological semantics of 3’UTR sequences. Specifically, to match the one-hot-coded matrix, we apply a linear projection that reduces the dimension of the unirna embeddings from 1024 to 32. The APARENT model is comprised of a shared model and two separate logistic regression layers, responsible for predicting isoform abundance and cleavage distribution respectively. The shared model is constructed from two convolutional layers as a feature encoder and two fully connected layers. The first convolutional layer has 96 convolutional kernels with a kernel size of 8. The second convolutional layer has 128 dimensional filters covering all 96 output channels from the previous layer, with a filter width of 6. BatchNorm layers and ReLU activation functions are applied after each convolutional layer, and a maxPool layer is applied after the first convolutional layer. The output of the shared model is then fed into two logistic regression layers, each with a bias weight vector indexed by the source UTR library, to derive the final prediction results.

#### Splice Site, RNA Modification, and Non-coding RNA Classes Prediction

We consider splice site, RNA modification and nc-RNA family classification as a sequence-level classification task. The CLS token of the sequence is fed into a two-layer fully connected neural network, and the cross-entropy loss function is used.

## 5 Acknowledgements

Patents have been filed based on the methods described in this manuscript.

